# Defective Neuronal Differentiation in Lowe Syndrome is Associated with Mitochondrial Dysfunction and Impaired Cilia-related Sonic Hedgehog Signaling

**DOI:** 10.1101/2024.11.01.621496

**Authors:** Grzegorz Walkiewicz, Siyu Chen, Chien-Hui Lo, Jingyu Zhao, Zhiquan Liu, Biao Wang, Benjamin Lawson, Qing Wang, Tia J. Kowal, Yang Sun

**Affiliations:** Department of Ophthalmology, Stanford University School of Medicine, Palo Alto, CA, USA; Palo Alto Veterans Administration, Palo Alto, CA, USA; Maternal Children Health Research Institute at Stanford, Stanford University School of Medicine, Palo Alto, CA, USA; BioX, Stanford University School of Medicine, Palo Alto, CA, USA

**Keywords:** Neuronal differentiation, Cilia formation, Mitochondria, ROS, Oxidative stress, Lowe Syndrome, OCRL

## Abstract

Human brain development requires tight coordination of metabolic and signaling pathways. Lowe syndrome (LS) is a recessive X-linked disorder characterized by proximal tubular renal disease, congenital cataracts, glaucoma, and neurodevelopmental delays. While LS results from mutations in the *OCRL* gene, which encodes an inositol polyphosphate 5-phosphatase, the cellular mechanisms driving neuronal dysfunction remain poorly understood. In this study, using patient-derived iPSC neurons, an *OCRL* knockout mouse model, and an independent zebrafish OCRL-deficient model, we identified mitochondrial dysfunction as a conserved phenotype of OCRL loss across species. Collectively, our findings showed that OCRL deficiency leads to reduced mitochondrial activity, decreased mtDNA levels, reduced mitochondrial content (TOM20), and increased oxidative stress. We further showed that OCRL-deficient neural cells exhibited an altered balance of neuronal versus astrocytic differentiation, rather than a defect in neurogenesis. Additionally, we observed impaired Sonic Hedgehog (Shh) signaling and ciliary homeostasis. Thus, we propose that mitochondrial dysfunction-induced oxidative stress acts as a central mediator linking OCRL loss to altered cell fate and disrupted Shh signaling, providing a unifying framework for these phenotypes.

## 1. Introduction

Lowe syndrome (LS) (OMIM #309000) is a rare X-linked disorder characterized by bilateral congenital cataracts, glaucomatous optic nerve degeneration, renal tubular dysfunction, and intellectual disability. The syndrome results from mutations in the oculocerebrorenal syndrome of Lowe gene (*OCRL*), which encodes an inositol polyphosphate 5-phosphatase that primarily hydrolyzes phosphatidylinositol 4,5-bisphosphate (PI(4,5)P2) to generate PI4P (Lewis et al., 1993; Luscher et al., 2019; Prosseda et al., 2017; Sakakibara et al., 2022; Sharma et al., 2015). Through its role in phosphoinositide metabolism, *OCRL* regulates key cellular processes including membrane trafficking, cytoskeletal organization, and intracellular signaling. Neurologically, patients with LS often exhibit developmental delays, intellectual disability, absent deep tendon reflexes, and hypotonia (Bökenkamp and Ludwig, 2016; Lewis et al., 1993). Nevertheless, the diagnosis of LS remains complicated due to the heterogeneity of its clinical presentation and overlap with other metabolic and neurodevelopmental disorders. This underscores the need for better-defined molecular mechanisms and potential biomarkers that could aid in diagnosis and disease stratification.

Several case reports point to the existence of a link between mitochondrial dysfunction and the pathogenesis of LS: (1) One report described a 5-year-old boy with *OCRL* mutation-confirmed LS who was initially diagnosed with mitochondriopathy based on electron microscopic evidence of mitochondrial abnormalities (Dumic et al., 2020). (2) Another patient, initially suspected of having chronic progressive external ophthalmoplegia (CPEO) due to mitochondrial disease, was later found to carry a missense mutation in *OCRL* (Craigen et al., 2013). Mitochondrial involvement in LS remains poorly defined, and the mechanisms linking *OCRL* function to mitochondrial homeostasis are not understood.

Increasing evidence demonstrates that reactive oxygen species (ROS) levels and mitochondrial DNA damage levels play an important role in directing neural stem cell (NSC) differentiation toward either neuronal or astroglial lineages (Adusumilli et al., 2021; Shahin et al., 2023; Wang et al., 2011). Previous studies have also established that astrocytes maintain their function independently of mitochondrial DNA integrity, while mitochondrial metabolism significantly contributes to astrocyte activation (Ignatenko et al., 2018; Wang et al., 2011). Astrocytes are essential for maintaining multiple critical functions in the central nervous system (CNS), including ionic balance, blood-brain barrier integrity, synaptic function, and metabolic homeostasis (Cabezas et al., 2014; McNeill et al., 2021; Oksanen et al., 2019; Pociūtė et al., 2024). In response to CNS injury, disease, or infection, astrocytes undergo a diverse array of morphological, molecular, and functional changes that are referred to as reactive astrogliosis (Escartin et al., 2021; Matusova et al., 2023; Zamanian et al., 2012). However, how metabolic stress, particularly mitochondrial dysfunction, modulates astrocyte behavior and neural lineage specification in neurodevelopmental disorders remains unclear. Based on these observations, we investigated the role of OCRL in mitochondrial dysfunction and increased oxidative stress in neuronal and astrocyte differentiation.

## 2. Results

### 2.1. Distinct differentiation of neuronal stem cells and neuronal progenitor cells (NSPCs) in *OCRL* knockout and LS iPSCs

To assess whether OCRL loss-of-function affects neuronal lineage specification, we employed an *in vitro* model of induced neurons (iNs) differentiated from Lowe syndrome (LS) patient-derived induced pluripotent stem cells (iPSCs) using a rapid single-step direct conversion protocol (Zhang et al., 2013). The LS iPSC line (LS100) was derived from a 17-year-old male patient clinically diagnosed with LS following genetic confirmation of an OCRL mutation. In contrast, the familial control line (LS200) originated from his 22-year-old unaffected brother (Barnes et al., 2018). Additionally, a CRISPR-Cas9-generated *OCRL* knockout (690 KO) iPSC line, previously generated using CRISPR-Cas9 from a healthy, unrelated control (690 Ctrl), was obtained from Herbert Lachman’s laboratory and used as an independent model of OCRL deficiency (Barnes et al., 2018). To verify pluripotency, all iPSC lines expressed canonical markers Nanog and Oct4, confirming robust stemness and proliferative capacity (Figure 1a, b). As expected, OCRL protein expression was absent in OCRL knockout iPSCs and markedly reduced in LS100 iPSCs compared to the WT-OCRL control line (LS 200) (Figure 1a, b). Following neuronal induction (Figure 1c), OCRL-deficient iPSCs exhibited altered differentiation patterns compared to control cells. Specifically, we observed an increased proportion of GFAP-positive cells, indicative of astrocytic identity, in OCRL-deficient cultures relative to controls (Figure 1d, e). To further characterize lineage specification, we performed qPCR analysis of neural markers. Expression of neuronal markers (*FOXG1, NEUN*) was higher in control and wild-type iNs (Figure 1f), whereas the astrocytic marker GFAP was significantly upregulated in OCRL knockout and LS-derived iNs (Figure 1g).

**Figure 1.**
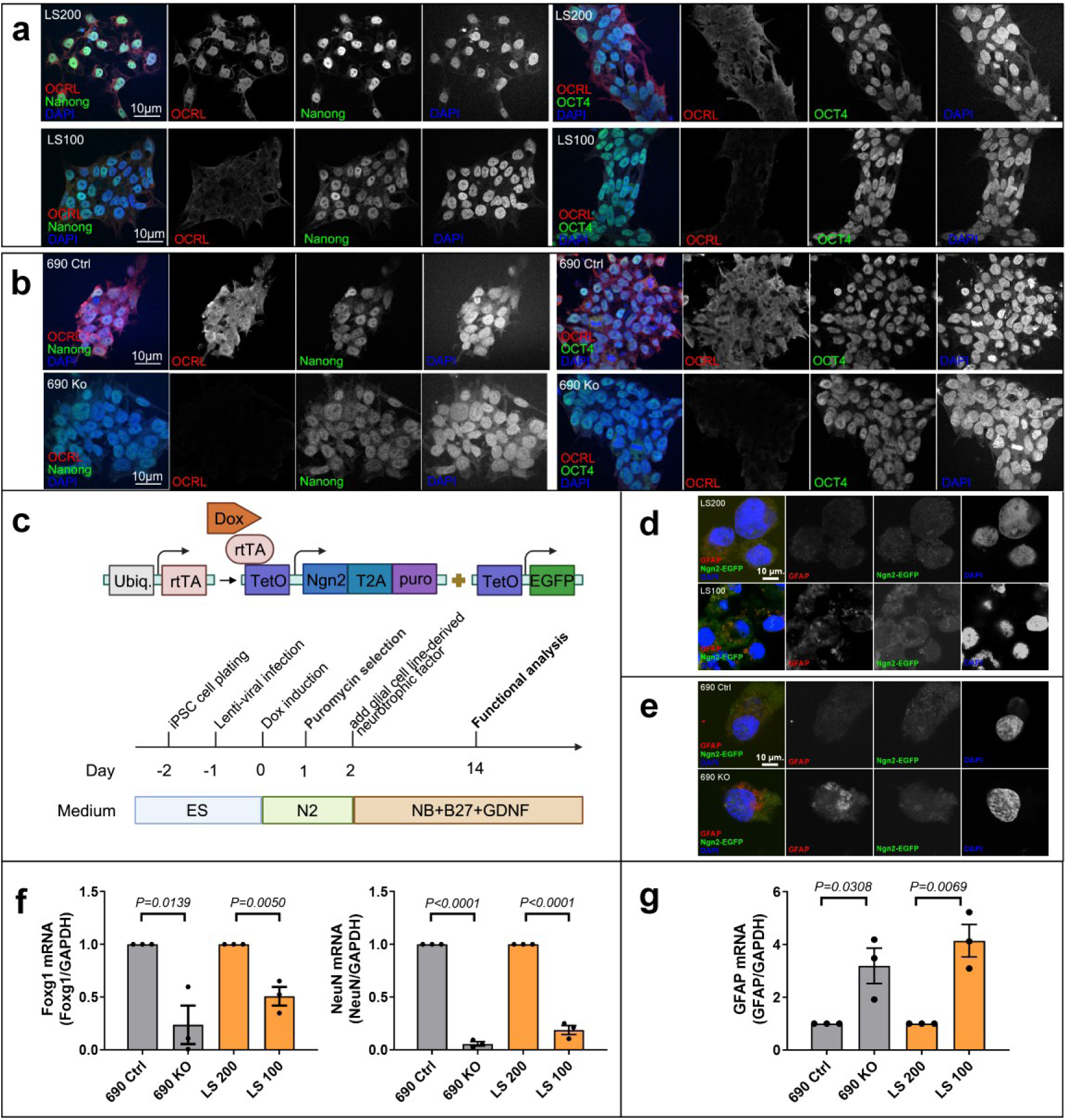
Increased astrocyte production during neuronal differentiation in OCRL-deficient and Lowe syndrome iPSCs. (a and b) Immunofluorescence analysis of OCRL expression and pluripotency markers in iPSCs. Cells were stained for OCRL (red) and pluripotency markers Nanog and OCT4 (green), with nuclei counterstained with DAPI (blue). Scale bars are as indicated. (c) Schematic representation of Ngn2-mediated direct conversion of iPSCs into induced neurons (iNs) using lentiviral vectors (adapted from Zhang et al., 2013). (d and e) Immunofluorescence analysis of iPSC-derived iNs following neuronal induction. Cells were stained for GFAP (red), with Ngn2-EGFP marking transduced cells. Nuclei were counterstained with DAPI (blue). Scale bars as indicated. (f) qPCR analysis of neuronal markers (*FOXG1* and *NEUN*) in iN cells derived from control and OCRL-deficient iPSCs. (g) qPCR analysis of *GFAP* expression in iN cells. Gene expression values were normalized to *GAPDH*. Data represent the mean ± SEM from three independent experiments. Statistical significance was determined using Student’s t-test. Changes in gene expression reflect relative marker levels and do not directly quantify cell-type proportions.

### 2.2. OCRL deficiency leads to mitochondrial dysfunction in OCRL knockout and LS-patient-derived iN cells

To investigate whether OCRL deficiency disrupts mitochondrial integrity and bioenergetic function during neuronal differentiation, we assessed mitochondrial activity in OCRL knockout and LS patient-derived iNs generated from iPSCs. Quantitative PCR analysis of mitochondrial DNA (mtDNA) genes *CO2* and *D-LOOP* revealed a marked reduction in mtDNA transcript levels in OCRL-deficient iNs compared to wild-type and sibling control iNs (Figure 2a). Consistently, immunostaining for 8-oxo-dG, a marker of oxidative DNA damage, demonstrated pronounced oxidative stress in OCRL knockout and LS iNs, whereas wild-type and control cells showed minimal reactivity (Figure 2b, c). Because these results suggested impaired mitochondrial oxidative phosphorylation (OXPHOS), we next evaluated mitochondrial respiratory function using Seahorse extracellular flux analysis. The oxygen consumption rate (OCR), a direct measure of OXPHOS efficiency, was significantly reduced in both *OCRL* knockout and LS-derived iNs compared with wild-type and unaffected control iNs (Figure 2d).

**Figure 2.**
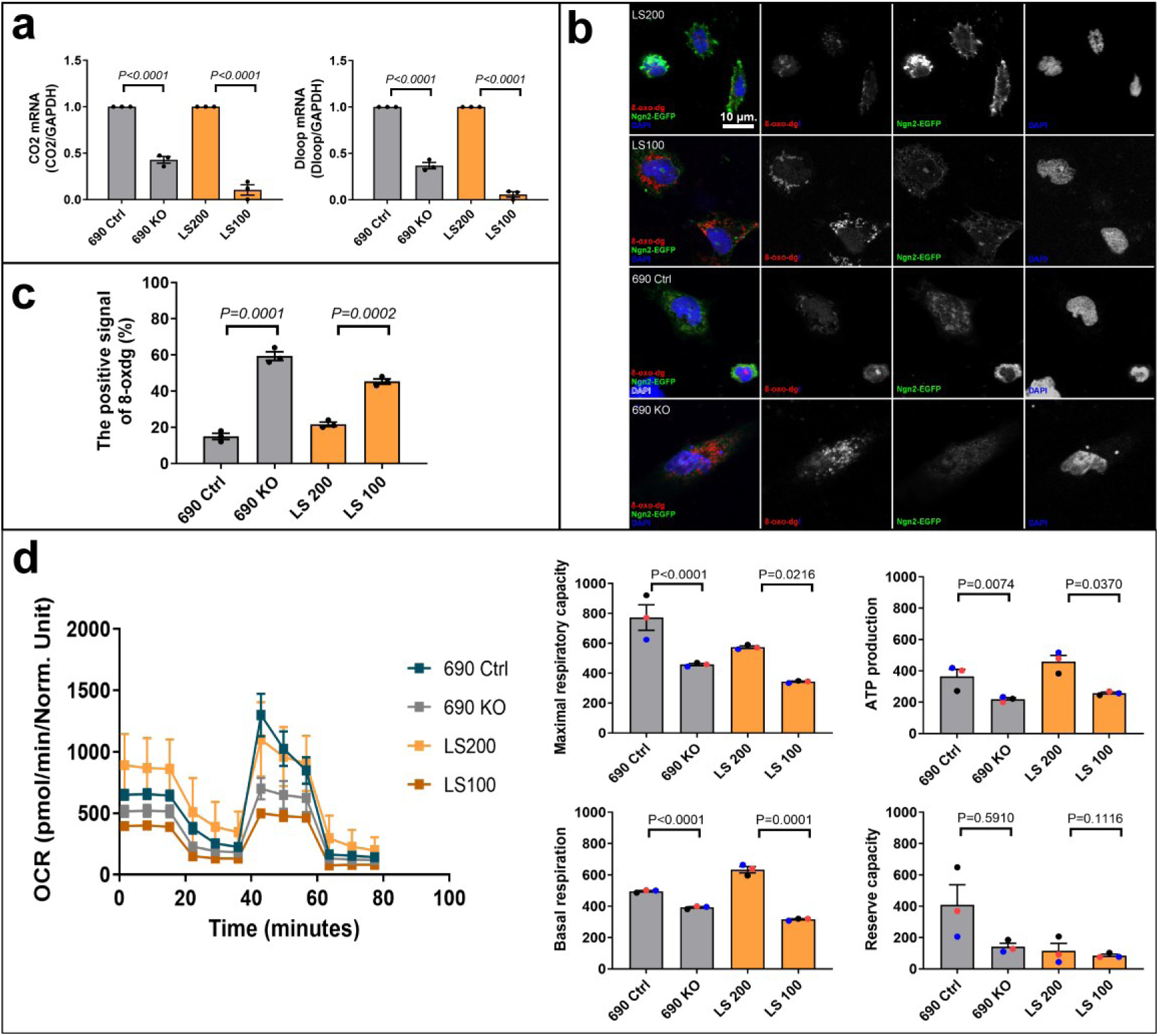
Altered mitochondrial parameters in OCRL-deficient iPSC-derived neurons. (a) qPCR analysis of mitochondrial DNA (mtDNA) levels, assessed using *CO2* and *D-loop* regions, in iN cells derived from control and OCRL-deficient iPSCs. (b) Immunofluorescence staining for 8-oxo-dG (red), a marker of oxidative DNA damage, in iN cells. Ngn2-EGFP (green) marks induced neurons. Nuclei are counterstained with DAPI (blue). Scale bars as indicated. (c) Quantification of the percentage of 8-oxo-dG-positive cells. More than 100 cells were analyzed per independent experiment. (d) Mitochondrial respiration was assessed by oxygen consumption rate (OCR) using Seahorse extracellular flux analysis. Gene expression values were normalized to GAPDH. Data represent mean ± SEM from three independent experiments. Statistical significance was determined using Student’s t-test.

### 2.3. Elevated astrocytic differentiation during neuronal differentiation of NSPCs in the Lowe syndrome (IOB) mouse model

Based on our iPSC-based findings, we hypothesized that CNS development in the LS mouse model would exhibit higher levels of astrocytic progenitor cells than neuronal progenitor cells. To examine neural cell composition *in vivo*, we analyzed brain tissue from the humanized Lowe syndrome (IOB) mouse model using adult mice (2-month-old mice), previously characterized for its ocular phenotype (Bothwell et al., 2011). Brain sections were assessed for the expression of neuronal and astrocytic markers (Supplementary Figure 3).

To further assess lineage specification, we quantified the expression of neuronal markers (*PAX6, NEUN*) (Figure 3a) and the astrocytic marker *GFAP* (Figure 3b) in the brain tissue. *GFAP* expression was increased in IOB brain tissue compared to wild-type controls, whereas expression of neuronal markers (*PAX6*, *NEUN*) was reduced (Figure 3a, b). Immunohistochemical analysis of brain sections showed an increased GFAP-positive signal relative to neuronal marker staining in IOB mice compared to controls (Figure 3c, d).

**Figure 3.**
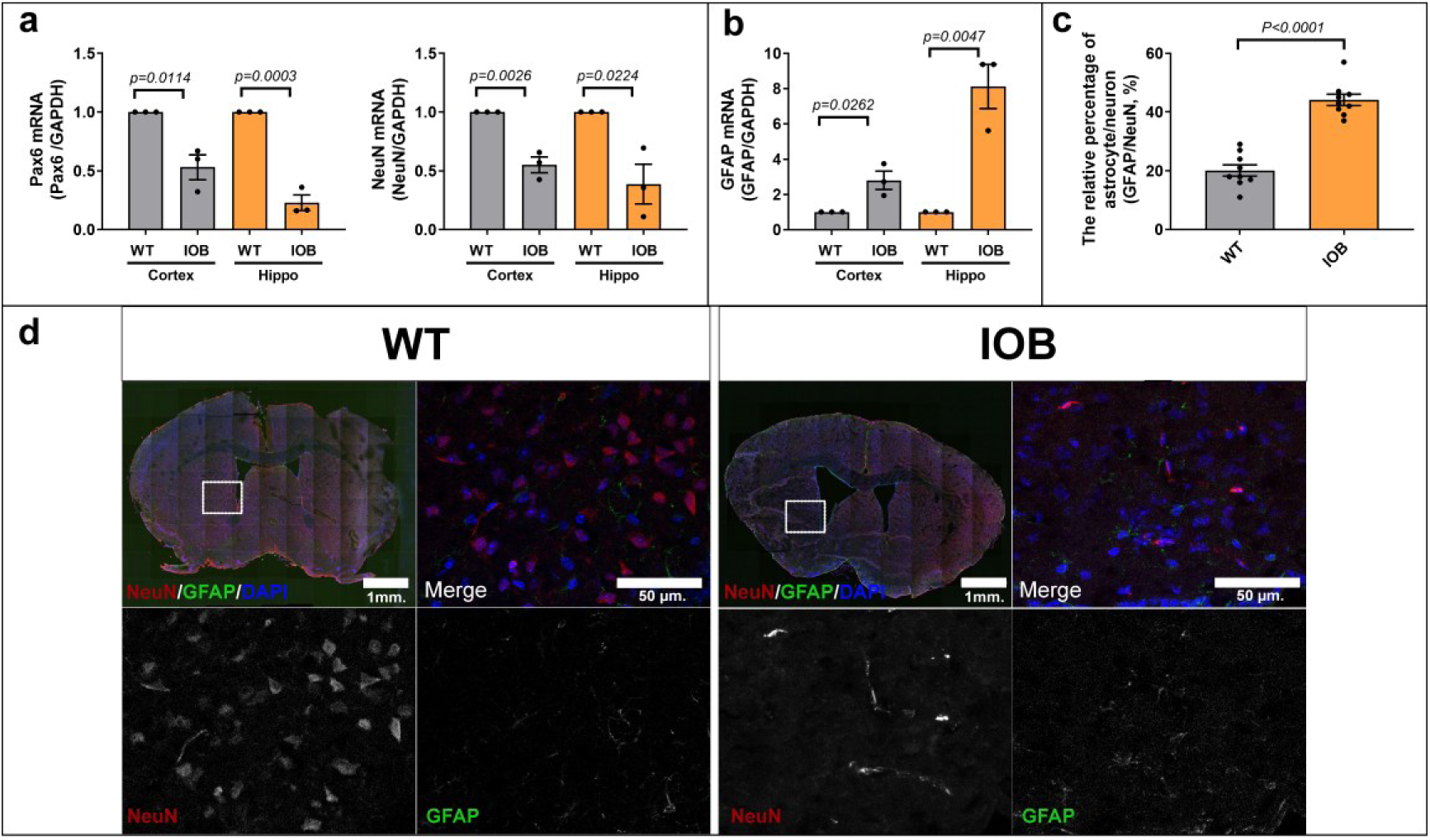
Altered neuronal and astrocytic marker expression in the Lowe syndrome mouse model. (a) qPCR analysis of progenitor-associated marker (*Pax6*) and neuronal markers (*NeuN*) in brain tissue. (b) qPCR analysis of astrocytic marker *GFAP* in brain tissue. (c) Quantification of NeuN and GFAP signal intensity in brain sections. More than 100 cells were analyzed per independent experiment. (d) Representative images of brains from wild-type (WT) and *INPP5B/OCRL* double knockout (IOB) mice. Immunofluorescence staining of brain sections for NeuN (red) and GFAP (green). Nuclei are counterstained with DAPI (blue). Scale bars as indicated. Gene expression values were normalized to GAPDH. Data represent mean ± SEM. Statistical significance was determined using Student’s t-test.

### 2.4. Reduced mitochondrial function during CNS development in the LS mouse model

To assess mitochondrial status *in vivo*, we analyzed mitochondrial DNA levels and oxidative stress in brain tissue from 2-month-old IOB mice and age-matched wild-type (WT) controls. Based on our findings from LS patient-derived iNs, we hypothesized that OCRL-deficiency *in vivo* similarly impairs mitochondrial function during CNS development. Quantitative PCR analysis revealed a significant reduction in mitochondrial DNA (mtDNA) measured by the *mito1 and COX1,* in IOB brain tissues compared with age-matched WT controls (Figure 4a). To evaluate mitochondrial oxidative stress, we performed immunostaining for 8-oxo-dG, a well-established marker of oxidative DNA damage (Hahm et al., 2022). IOB brain sections displayed an increased 8-oxo-dG-positive signal compared to WT controls (Figure 4b, c).

**Figure 4.**
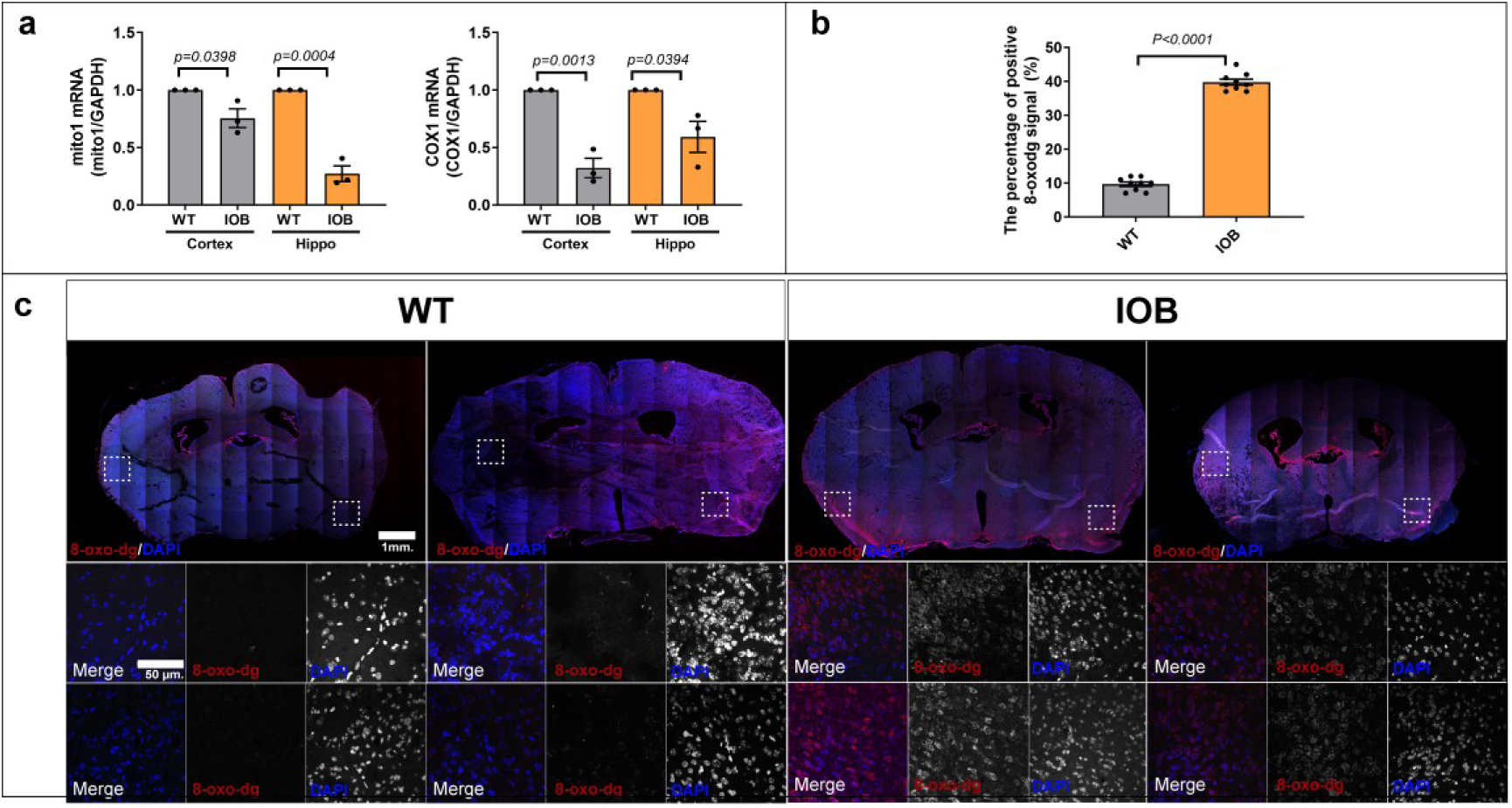
Altered mitochondrial parameters in the Lowe syndrome mouse brain. (a) qPCR analysis of mitochondrial DNA (mtDNA), assessed using *mito1* and *COX1* in brain tissue from WT and IOB mice. (b) Quantification of 8-oxo-dG-positive signal in brain sections. More than 100 cells were analyzed per independent experiment. (c) Immunofluorescence staining for 8-oxo-dG (red) in brain sections. Nuclei are counterstained with DAPI (blue). Scale bars as indicated. Gene expression values were normalized to GAPDH. Data represent mean ± SEM. Statistical significance was determined using Student’s t-test.

### 2.5. Altered mitochondrial parameters in OCRL-deficient zebrafish larvae

To assess mitochondrial parameters in an independent *in vivo* model, we analyzed OCRL-deficient zebrafish larvae generated by CRISPR/Cas9-mediated targeting of the zebrafish *ocrl* gene, the homolog of human *OCRL* (Supplementary Figure 4). OCRL-deficient larvae exhibited developmental abnormalities compared to control gRNA-injected larvae (Figure 5a) and showed reduced survival over time (Figure 5b,c). Mitochondrial reactive oxygen species (ROS) were evaluated using MitoSOX staining. OCRL-deficient larvae displayed increased MitoSOX signal in both cranial and ocular regions compared to controls (Figure 5d). Mitochondrial content was assessed by TOM20 immunostaining. OCRL-deficient larvae showed reduced TOM20 signal relative to control larvae (Figure 5e). Mitochondrial membrane potential (ΔΨm) was measured using MitoTracker CMXRos. OCRL-deficient larvae exhibited reduced MitoTracker CMXRos intensity in cranial and ocular regions compared to controls (Figure 5f).

**Figure 5.**
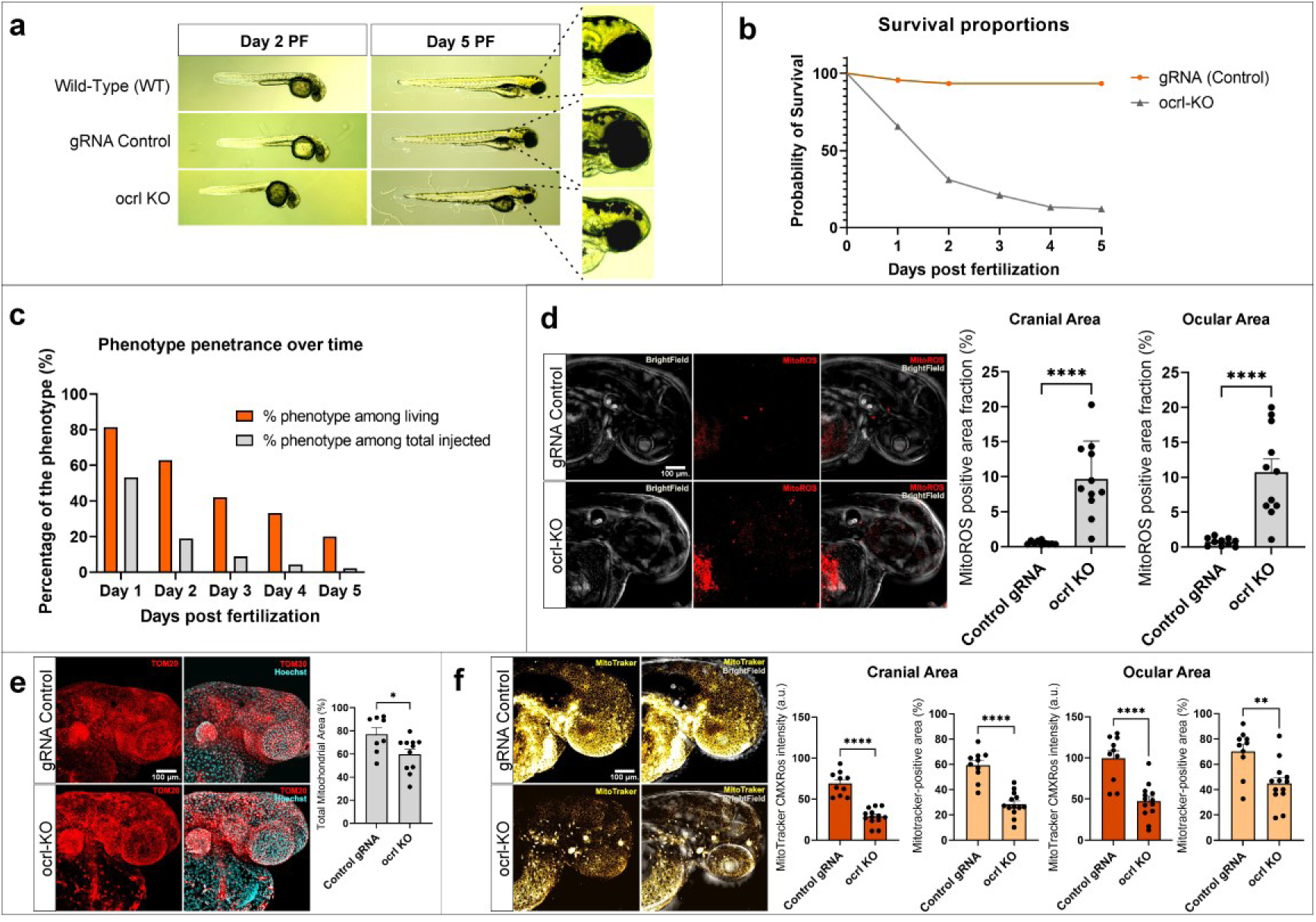
OCRL loss is associated with mitochondrial dysfunction, oxidative stress, and reduced survival in zebrafish. (a) Representative brightfield images of zebrafish larvae at 2 and 5 days post-fertilization (dpf), including wild-type (WT), control gRNA-injected, and *ocrl* knockout (KO) groups. (b) Kaplan-Meier survival analysis of zebrafish larvae. OCRL-deficient larvae exhibit reduced survival compared to control gRNA-injected larvae. (c) Quantification of phenotype penetrance over time (1-5 dpf), presented as (i) percentage of affected larvae among living animals and (ii) percentage of affected larvae relative to total injected embryos. (d) Mitochondrial reactive oxygen species (ROS) assessed by MitoSOX staining. Representative images and quantification of MitoSOX-positive area fraction (%) in cranial and ocular regions are shown. (e) Mitochondrial content assessed by TOM20 immunostaining. Representative images and quantification of TOM20-positive area fraction (%) are shown. (f) Mitochondrial membrane potential (ΔΨm) assessed by MitoTracker CMXRos staining. Representative images and quantification of MitoTracker CMXRos intensity (a.u.) and positive area fraction (%) in cranial and ocular regions are shown. Data are presented as mean ± SEM from n = 10-15 larvae per group. Statistical significance was determined using Student’s t-test unless otherwise indicated. Imaging and quantification were performed under identical conditions across all groups.

### 2.6. Defective ciliary homeostasis underlies neuronal dysfunction in Lowe syndrome

Given the mitochondrial dysfunction observed in OCRL-deficient models, we hypothesized that defective mitochondrial function may secondarily impair ciliary structure and Shh signaling in LS. To assess Shh pathway activity, we analyzed gene expression in iNs derived from OCRL knockout and LS patient-derived iPSCs. Quantitative PCR analysis revealed a marked downregulation of *GLI1, PTCH1,* and *Shh* transcripts in OCRL-deficient iNs compared with wild-type and unaffected sibling control iNs (Figure 6a). Consistently, primary cilia were examined in brain sections from IOB mice. IOB brain sections exhibited a significant reduction in the number of ciliated cells and elongated cilia length compared to wild-type controls (Figure 6b, c). To further assess the impact on Shh signaling *in vivo*, we quantified the mRNA levels of key pathway components, including *Gli1, Gli2, Gli3*, and *Ptch1*, in IOB and WT brains. All four transcripts were significantly reduced in IOB mice (Figure 6d). Western blot analysis confirmed decreased protein expression of *Shh* and *Gli1* in IOB brains compared to WT (Figure 6e).

**Figure 6.**
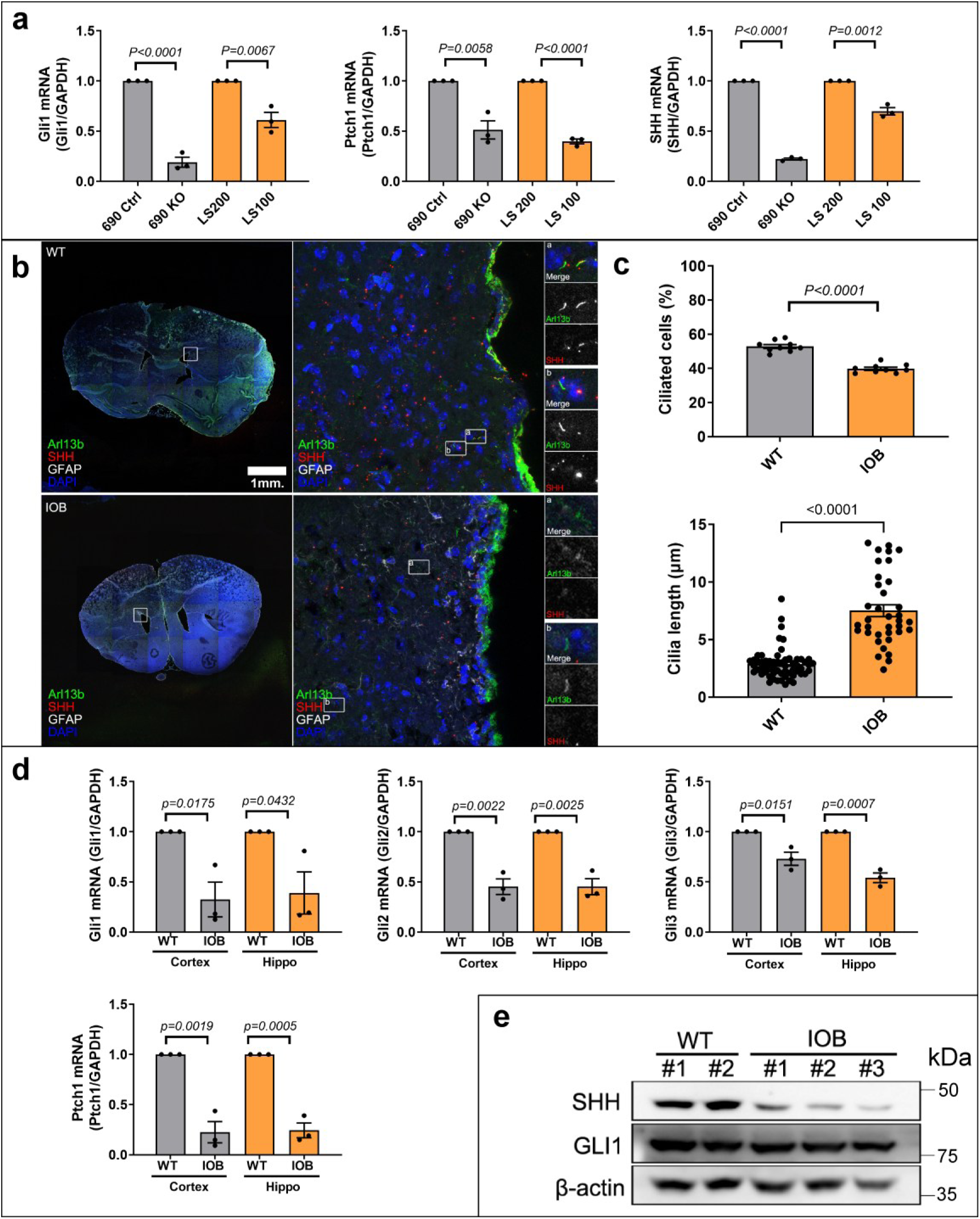
Altered ciliary parameters and Sonic Hedgehog signaling in OCRL-deficient models. (a) qPCR analysis of Shh pathway genes (*GLI1, PTCH1, SHH*) in iPSC-derived iN cells. (b) Immunofluorescence staining of brain sections for SHH (red) and the ciliary marker ARL13B (green). Nuclei are counterstained with DAPI (blue). Scale bars as indicated. (c) Quantification of the ciliated cells and cilia length in brain sections. More than 100 cells were analyzed per independent experiment. (d) qPCR analysis of Hedgehog pathway genes (*Gli1, Gli2, Gli3, Ptch1*) in brain tissue from WT and IOB mice. (e) Western blot analysis of SHH and GLI1 protein levels in brain tissue. β-actin was used as a loading control. Gene expression values were normalized to GAPDH. Data represent mean ± SEM. Statistical significance was determined using Student’s t-test.

## 3. Discussion

In this study, we expand our analysis across multiple model systems, including patient-derived iPSCs, CRISPR knockout human cells, the IOB mouse model, and an independent zebrafish OCRL-deficient model, to identify mitochondrial dysfunction as a conserved phenotype of OCRL loss. Across all models, OCRL loss was consistently associated with reduced mitochondrial DNA levels, decreased oxidative phosphorylation, and increased oxidative stress. In zebrafish, these alterations were further supported by reduced mitochondrial membrane potential, increased mitochondrial ROS, and decreased TOM20 staining, indicating reductions in both mitochondrial function and mitochondrial content. Together, these findings support the role of mitochondrial dysfunction as a contributing cause for OCRL deficiency in Lowe syndrome. Consistent with this, recent studies independently reported mitochondrial dysfunction alongside altered neurodevelopmental trajectories in OCRL-deficient systems, including enhanced Notch-dependent gliogenesis and impaired mitochondrial structure and function (Philip et al., n.d.; Sharma et al., 2025). Collectively, these studies support that mitochondrial dysregulation is a conserved feature of neurologic abnormalities in Lowe syndrome. These mitochondrial defects coincided with a consistent shift in lineage specification. Our data showed an altered balance between neuronal and astrocytic differentiation. This distinction supports that OCRL deficiency influences lineage specification rather than completely blocking neuronal development. In line with our observations, Sharma et al. recently showed enhanced Notch-dependent glial differentiation and delayed neuronal maturation in OCRL-deficient models (Sharma et al., 2025). Together, these studies support the concept that OCRL deficiency alters cell-fate decisions during neural development. Importantly, our data do not directly address Notch signaling, but instead identify mitochondrial dysfunction and altered ciliary/Shh signaling as additional pathways associated with this phenotype.

In addition to mitochondrial abnormalities, we identified alterations in ciliary homeostasis and Shh signaling. OCRL-deficient models exhibited reduced expression of key Shh pathway components at both the transcript and protein levels. In the IOB mouse brain, this was accompanied by a decreased number of ciliated cells together with increased cilia length, indicating altered ciliary homeostasis. Given the central role of the primary cilium in Shh signal transduction, these findings implicate disrupted ciliary/Shh signaling as an additional feature of OCRL deficiency (Zhuang, 2025). Notably, mitochondrial imbalance was shown to alter cilia length and disrupt cilia-dependent processes *in vivo*, linking bioenergetic state to developmental patterning (Burkhalter et al., 2019; Moruzzi et al., 2022).

Previous studies have reported adaptive ciliary responses to mitochondrial stress, including increased ciliogenesis and ciliary remodeling driven by ROS signaling and transcriptional activation of ciliogenic programs (Ignatenko et al., 2023; Moruzzi et al., 2022). In contrast, OCRL-deficient models in our study exhibited a reduced number of ciliated cells together with altered ciliary morphology, suggesting impaired ciliary homeostasis rather than a compensatory ciliogenic response. This difference may reflect the nature of OCRL deficiency, which represents a chronic disruption of phosphoinositide metabolism and endosomal trafficking rather than an acute mitochondrial stress response (Mehta et al., 2014). Prior studies established zebrafish as a relevant model for Lowe syndrome, revealing defects in neuroepithelial development and endocytic trafficking, including seizure-like phenotypes (Williams et al., 2022). Our findings extend these observations by linking these developmental abnormalities to mitochondrial dysfunction as a potential underlying mechanism. Given the critical role of OCRL in membrane dynamics at the ciliary base, its loss is likely to impair the formation and maintenance of cilia despite the presence of mitochondrial stress signals (Luo et al., 2012). Sustained mitochondrial dysfunction may therefore be associated with defective ciliary signaling rather than adaptive ciliogenesis (Kim et al., 2025; Moruzzi et al., 2022).

Thus, we propose a model in which OCRL deficiency leads to mitochondrial dysfunction and increased oxidative stress, which are associated with alterations in neural lineage balance and disruption of Shh signaling (Figure 7). While the causal relationships between these phenotypes remain to be established, oxidative stress may represent a potential link between mitochondrial impairment and downstream developmental signaling pathways. Shh signaling is a key regulator of neural progenitor fate and astrocyte development and contributes to the maintenance of region-specific astrocyte functions in the central nervous system (Garcia, 2021; Gingrich et al., 2022; Hill et al., 2021, 2019). Given the established roles of astrocytes in synapse formation, maturation, and elimination (Akdemir et al., 2020; Clavreul et al., 2022; Farhy-Tselnicker and Allen, 2018), disruption of these pathways may contribute to the altered neuron-to-astrocyte balance observed in OCRL-deficient systems. Importantly, our findings do not exclude the Notch-dependent mechanism described by Sharma et al. (Sharma et al., 2025), but suggest that mitochondrial dysfunction and altered ciliary/Shh signaling represent additional pathways associated with neurodevelopmental abnormalities in Lowe syndrome.

**Figure 7.**
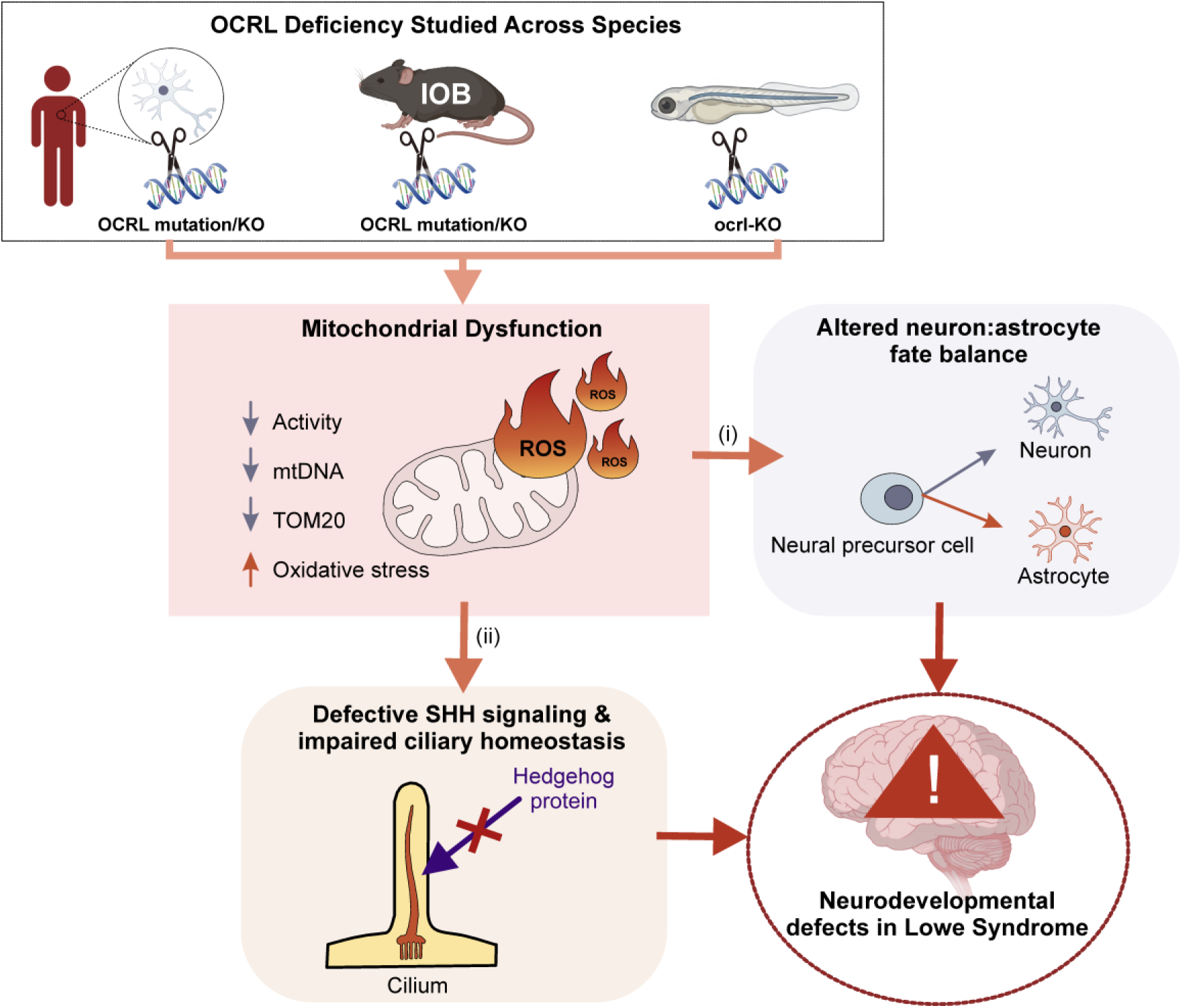
OCRL deficiency disrupts neuronal development through mitochondrial dysfunction, oxidative stress, and impaired Shh-cilia signaling. Schematic representation of the integrated findings across experimental systems. OCRL deficiency leads to mitochondrial dysfunction, characterized by reduced mitochondrial DNA, decreased oxidative phosphorylation, reduced mitochondrial content, and increased oxidative stress. Elevated oxidative stress is associated with two parallel processes: (i) altered balance between neuronal and astrocytic cell states and (ii) reduced Shh signaling, accompanied by changes in ciliary parameters, including decreased proportion of ciliated cells and increased cilia length. These combined alterations are associated with impaired neuronal development in Lowe syndrome.

Several limitations should be considered. The *in vivo* analyses were limited to available brain regions, and astrocytic differentiation was primarily assessed using GFAP expression. Furthermore, while mitochondrial dysfunction, altered neural differentiation, and impaired ciliary/Shh signaling were consistently observed across models, the causal relationships between these phenotypes remain to be determined.

In summary, our findings identify mitochondrial dysfunction and altered ciliary signaling as key features of OCRL deficiency across multiple model systems. By integrating these observations, we propose a conserved mitochondria-ROS-signaling axis associated with altered neural differentiation in Lowe syndrome (Figure 7). This framework highlights mitochondrial homeostasis and ciliary signaling as interconnected processes in neurodevelopment and suggests that targeting bioenergetic or redox pathways may represent a potential therapeutic avenue in Lowe syndrome.

## 4. Materials and methods

### 4.1. iPSCs culture and reagent

iPSCs were cultured on matrigel (Corning, 354277) in mTeSR1 Plus medium (Stem Cell Technologies, 85850). Media were changed daily, and confluent cells were passaged (1:2) using ReLeSR (Stem Cell Technologies, 05872). All cells were maintained at 37° C, 5% CO_2_.

### 4.2. Animals

All animal experiments adhered to the guidelines of the Association for Research in Vision and Ophthalmology Statement for the Use of Animals in Ophthalmic and Vision Research and were approved by the Institutional Animal Care and Use Committee (IACUC) of Stanford University School of Medicine.

#### 4.2.1. Mice

2-month-old Ocrl^−/−^ Inpp5b^−/−^ INPP5B^+/+^ (IOB) mice were generously provided by Robert L. Nussbaum (University of California, San Francisco). Wild-type (C57BL/6) mice from the Jackson Laboratories were used as controls for the IOB mice. The animals were housed under a 12-hour light/dark cycle with free access to water and food. Mice were anesthetized with isoflurane, with oxygen flow set to 2 liters per minute and isoflurane at 1% delivered via a nose cone.

#### 4.2.2. Zebrafish husbandry and generation of OCRL-deficient zebrafish larvae

Wild-type zebrafish (Danio rerio) were maintained under standard conditions at 28.5 °C on a 14 h light / 10 h dark cycle. Embryos were obtained by natural mating and raised in E3 embryo medium. All procedures were conducted in accordance with institutional animal care guidelines and approved protocols.

OCRL-deficient zebrafish were generated using CRISPR/Cas9-mediated gene disruption targeting the zebrafish *ocrl* ortholog. The zebrafish experimental procedures were adapted from a previously established protocol (Pagán et al., 2022). Briefly, guide RNAs (gRNAs) were designed to target coding regions of *ocrl* using CRISPR design tools. Synthetic crRNA and tracrRNA (IDT) were complexed with recombinant Cas9 protein (IDT, Alt-R Cas9 V3) to form ribonucleoprotein (RNP) complexes. The RNP complex was prepared as follows: the crRNA:tracrRNA duplex was annealed at 95°C for 5 minutes and cooled to room temperature, then incubated with Cas9 protein for 10-15 minutes at room temperature. Approximately 1-2 nL of RNP complex was microinjected into one-cell-stage zebrafish embryos using a microinjection system. As a control, embryos were injected with a non-targeting gRNA (out-of-genome control) to account for CRISPR-associated toxicity. Injected embryos were maintained at 28.5 °C and screened for phenotypic abnormalities and survival. Larval survival was monitored daily from 1 to 5 days post-fertilization (dpf). Dead larvae were removed at each time point. Developmental phenotypes were assessed by brightfield imaging using a stereomicroscope, with particular attention to body morphology, cranial structure, and eye development.

### 4.3. Zebrafish mitochondrial assays (MitoSOX, MitoTracker CMXRos, and TOM20 staining)

Mitochondrial function, reactive oxygen species (ROS), and mitochondrial content were assessed in OCRL-deficient zebrafish larvae at 2 days post-fertilization (2 dpf). To assess mitochondrial membrane potential (ΔΨm) and mitochondrial ROS, live zebrafish larvae were stained with MitoTracker™ Red CMXRos (Thermo Fisher Scientific) and MitoSOX™ Red (Thermo Fisher Scientific). Larvae at 2 dpf were incubated in E3 medium containing 100 nM MitoTracker CMXRos and 2 µM MitoSOX for 20 minutes at 28.5 °C in the dark. Following incubation, larvae were washed twice in fresh E3 medium to remove excess dye, anesthetized using tricaine (MS-222), and mounted in low-melting-point agarose in a confocal imaging dish. Fluorescent images were acquired using an Olympus fluorescence microscopy system under identical acquisition settings across experimental groups. Quantification was performed using Fiji/ImageJ by measuring mean fluorescence intensity or positive area fraction (%) within defined regions of interest (ROIs) corresponding to cranial and ocular regions. For each larva, cranial and ocular regions were analyzed as predefined ROIs. Mitochondrial content was quantified as TOM20-positive area fraction (%) using Fiji/ImageJ. N = 10-15 larvae per group were analyzed.

### 4.4. Plasmids

The lentiviral vectors for Ngn2-mediated conversion of iPSCs to iN cells are from Thomas C. Sudhof’s lab (Zhang et al., 2013).

### 4.5. Lentivirus production and infection

5×10^5^ 293FT cells were plated on 60-mm dishes using TurboFect™ Transfection Reagent with the following plasmids: 1.5 μg of V-SVG, 2.5 μg of pCMV-gag-pol, and 3.5 μg of the lentiviral vector DNA constructs. The supernatant containing viral particles was harvested 48 h after transfection. Virus-containing media was passed through a 0.45μm filter (Fisher Scientific, 13-100-105).

### 4.6. Generation of iN Cells from Human iPSCs

iN cells were generated using an established Ngn2-based direct conversion protocol (Zhang et al., 2013). iPSCs were treated with Accutase (Stem Cell Technologies, 07920) and plated as dissociated cells in 24-well plates (iPSCs: 1.5 x 10^4^ cells/well) on day 2 (Figure 1c). Cells were plated on matrigel (Corning, 354277)-coated coverslips in mTeSR1 medium. On day 1, lentivirus prepared as described above (0.3 ml/well of 24-well plate) was added to fresh mTeSR1 medium containing polybrene (8 mg/mL, Sigma). On day 0, the culture medium was replaced with DMEM/F12 (Thermo Fisher Scientific, 11-330-057) containing N2(STEMCELL Technologies, 07152), NEAA (Thermo Fisher Scientific, 11-140-050), human BDNF (10 mg/L, STEMCELL Technologies, 78058), human NT-3 (10 mg/L, PeproTech, 450-03), and mouse laminin (0.2 mg/L, Thermo Fisher Scientific, 23017015). Doxycycline (2 g/L, Fisher Scientific, AC446060050) was added on day 0 to induce TetO gene expression and retained in the medium until the end of the experiment. On day 1, a 24 hr puromycin selection (1 mg/L) period was started. On day 2, replace with Neurobasal medium (Thermo Fisher Scientific, 21103049) supplemented with B27/Glutamax (Invitrogen) containing BDNF and NT3. After day 2, 50% of the medium in each well was exchanged every 2 days. FBS (2.5%) was added to the culture medium on day 10 to support astrocyte viability, and iN cells were assayed on day 14 or 21 in most experiments.

### 4.7. Immunostaining

#### 4.7.1. Cell immunostaining

Cells were cultured on coverslips coated with 0.1 mg/mL poly-L-lysine and fixed with methanol at -20°C for 15 minutes. The cells were then washed three times with PBS and incubated in a blocking buffer containing 3% bovine serum albumin (w/v) and 0.1% Triton X-100 in PBS for 30 minutes at room temperature (RT). Primary antibodies, diluted in the blocking buffer, were applied for 2 hours at RT. Alexa Fluor 488-, 594-, or 647-conjugated goat secondary antibodies (Thermo Fisher Scientific) were used at a 1:500 dilution and incubated for 1 hour at RT. DNA was stained with 4,6-diamidino-2-phenylindole (DAPI; Thermo Fisher Scientific). Coverslips were then mounted on slides using ProLong™ Gold Antifade mounting medium (Thermo Fisher Scientific).

#### 4.7.2. Brain section immunostaining

Mouse brains were harvested and fixed in 4% paraformaldehyde (PFA) at 4°C, followed by cryoprotection in sucrose solution (e.g., 30% sucrose in PBS). Brains were embedded in optimal cutting temperature (OCT) compound and sectioned using a cryostat to obtain 20 µm sections. Brain sections were permeabilized in 0.3-0.5% Triton X-100 in PBS for 20-30 minutes and blocked in 5% BSA and 0.1% Triton X-100 in PBS for 1 hour at RT. Sections were incubated with primary antibodies diluted in blocking buffer overnight at 4°C. After washing, sections were incubated with appropriate Alexa Fluor-conjugated secondary antibodies (1:500) for 1 hour at RT. Nuclei were stained with DAPI. Sections were mounted using an antifade mounting medium and imaged using a confocal microscope under identical acquisition settings across samples.

#### 4.7.3. Zebrafish whole-mount immunostaining

For whole-mount staining, zebrafish larvae at 2 dpf were fixed in 4% PFA overnight at 4°C, washed in PBS, and permeabilized with 0.5% Triton X-100 in PBS for 30 minutes. Blocking was performed in 5% BSA and 0.1% Triton X-100 in PBS for 1 hour at RT. Larvae were incubated with primary antibodies (e.g., anti-TOM20, 1:200) overnight at 4°C, followed by washing and incubation with Alexa Fluor-conjugated secondary antibodies (1:500) for 1 hour at RT. Samples were mounted in low-melting-point agarose for imaging.

#### 4.7.4. Imaging

Fluorescent images were acquired using a Zeiss LSM880 confocal microscope or an Olympus fluorescence microscope, depending on the experimental setup. All images were acquired using identical settings across experimental groups. Image processing and quantification were performed using ZEN software (Carl Zeiss) or Fiji/ImageJ (NIH)

### 4.8. Immunoblotting

Cells were washed twice with ice-cold PBS and lysed in ice-cold RIPA lysis buffer (Millipore, 20-188) containing a protease inhibitor cocktail (Thermo Fisher Scientific, PI78430). The lysate was centrifuged at 13,500 g for 15 minutes at 4°C to remove cell debris. Protein concentrations were measured using the BCA Protein Assay (Thermo Fisher Scientific, 23227). Equal amounts of protein were combined with SDS sample buffer, boiled at 95°C for 5 minutes, and separated by SDS-PAGE. The proteins were then transferred to 0.2 µm nitrocellulose membranes (Bio-Rad, 1620097). The membranes were blocked for 1 hour at room temperature (RT) with 5% non-fat milk in TBS-T (20 mM Tris, pH 7.6, 137 mM NaCl, and 0.1% Tween-20) and incubated overnight at 4°C with primary antibodies in the blocking solution. The membranes were washed three times with TBS-T and incubated with HRP-conjugated anti-mouse or anti-rabbit secondary antibodies (Invitrogen, 31430 and 31460) for 1 hour at RT. After three additional washes with TBS-T, the proteins were visualized using ECL Western blotting substrate (Thermo Fisher Scientific, 34095).

### 4.9. Primary antibodies

Primary antibodies were obtained from the following sources and used according to the manufacturers’ instructions: mouse IgG1 anti-OCRL/INPP5b, NeuroMab clone N166A/26 (IF 1: 250; UC Davis/NIH NeuroMab Facility), rabbit anti-Nanog (IF 1: 250; 3580S, Cell Signaling Technology), rabbit anti-Oct-4A (C30A3) (IF 1: 250; 2840S, Cell Signaling Technology), Chicken anti-GFAP(IF 1: 250; ab4674, Abcam), mouse IgG2b anti-8-oxo-Dg (IF 1: 250; 4354-MC-050; R&D systems), rabbit anti-Sonic Hedgehog antibody [EP1190Y] (IF: 1:200; WB: 1:500: ab53281, Abcam), rabbit anti-Gli1 antibody (WB: 1:500: ab217326, Abcam), mouse anti-Arl13b antibody (N295B/66) (IF: 1:500; 75-287, Antibodies Incorporated), mouse anti-NeuN Antibody, clone A60 (IF: 1:200; MAB377, Sigma-Aldrich), mouse anti-β actin (WB: 1:5000: 66009-1, Proteintech), mouse anti-TOM20 (IF 1:200; ab56783, Abcam)

### 4.10. Quantitative real-time PCR (qPCR)

Quantitative real-time PCR (qPCR) was conducted using HiScript III RT SuperMix for qPCR plus gDNA wiper (Vazyme, R323-01). qPCR was performed using FastSYBR Mixture (2X) (CWBio, CW0955L). The amplification was carried out in 20 μl reaction mixtures containing 100 ng of total DNA, 1X SYBR-Green PCR Master Mix, and 0.5 μM of each primer. Each marker was tested in triplicate reactions in a 96-well plate using a three-step amplification protocol: initial denaturation at 95°C for 5 minutes, followed by 40 cycles of 95°C for 15 seconds, 60°C for 30 seconds, and 72°C for 30 seconds. Relative gene expression was calculated using the comparative Ct (ΔΔCt) method and normalized to GAPDH as an internal control. Each sample was analyzed in triplicate, and no-template controls were included. The sequences of each gene were shown in Supplemental Figures 1 and 2.

### 4.11. Oxygen consumption rate (OCR)

Oxygen consumption rate (OCR) was measured using a Seahorse Biosciences XFe96 extracellular flux analyzer. Cells were seeded at a density of 1.25 × 10^5^ cells per well in XFe96 cell culture plates. After 24 hours, cell attachment was confirmed, and the cells were incubated overnight at 37°C with 5% CO2. Prior to the assay, the cells were switched to Seahorse XF DMEM medium containing 1 mM pyruvate, 2 mM glutamine, and 10 mM glucose and equilibrated for 1 hour at 37°C without CO2. OCR was then measured using the following inhibitors: 2.5 μM oligomycin, 2 μM carbonyl cyanide 4-(trifluoromethoxy) phenylhydrazone (FCCP), and 0.5 μM rotenone combined with 0.5 μM antimycin A (Agilent Technologies, 103015-100). Each condition was tested in triplicate cycles, consisting of 3 minutes of mixing followed by 3 minutes of measurement. After the assay, the cell number per well was determined using the Cytation 5, and the OCR was normalized to the cell number for each well.

### 4.12. Primary cilia quantification

Primary cilia were visualized by immunostaining using ARL13B as a ciliary marker. Images were acquired using a confocal microscope under identical acquisition settings across all samples. The proportion of ciliated cells was quantified by manually counting ARL13B-positive cilia relative to total nuclei (DAPI) within defined regions of interest (ROIs). Cilia length was measured manually using Fiji/ImageJ. For each cilium, a line was drawn along the length of the ARL13B-positive structure using the segmented line tool, and the length was recorded in micrometers (µm) after spatial calibration. All measurements were performed using identical thresholding and analysis settings across groups.

### 4.13. Statistical data analysis

All data are presented as the mean with standard deviation (SD) from at least 3 independent experiments. Experimental samples and numbers for statistical testing are reported in the corresponding figure legends. All p-values are from Student’s t-tests for two-group comparisons (GraphPad Prism 8).

## 5. Conflict of Interest Statement

The authors declare no competing interests.

## 6. Author Contributions

GW, SC, and C-HL designed and carried out the experiments, data analysis, and wrote the manuscript. SC, ZL, JZ, and BL contributed to the manuscript editing and experiment assistance. TK and BW contributed to setting up iPSCs processing. BW and QW contributed to setting up the IOB mouse processing. YS supervised the project.

## 7. Acknowledgements

We thank Dr. Robert L. Nussbaum (University of California, San Francisco) for generously sharing the IOB mouse strain. We thank Dr. Herbert M. Lachman’s lab for the iPSCs of LS.

## 8. Ethics Statement

All animal experiments followed the guidelines of the Association for Research in Vision and Ophthalmology Statement for the Use of Animals in Ophthalmic and Vision Research and were approved by the Institutional Animal Care and Use Committee (IACUC) of Stanford University School of Medicine.

## 9. Funding Statement

This work was supported by RO1-EY32159 (YS), EY-034932 (YS), R01-EY025295 (Y.S.), and an unrestricted grant from Research to Prevent Blindness, New York, NY. VA merit CX001481 (Y.S.), Ziegler Foundation for the Blind (Y.S.), and Children’s Health Research Institute Award (Y.S.). Research for Prevention of Blindness Unrestricted grant and NIH P30EY026877 (Stanford Ophthalmology) and R38EY037090 (Stanford Ophthalmology).

## Notes

### Competing Interest Statement

The authors have declared no competing interest.

### Summary of Updates

Lowe syndrome (LS) is a recessive X-linked disorder characterized by proximal tubular renal disease, congenital cataracts, glaucoma, and neurodevelopmental delays. While LS results from mutations in the OCRL gene, which encodes an inositol polyphosphate 5-phosphatase, the cellular mechanisms driving neuronal dysfunction remain poorly understood. In this study, using patient-derived iPSC neurons, an OCRL knockout mouse model, and an independent zebrafish OCRL-deficient model, we identified mitochondrial dysfunction as a conserved phenotype of OCRL loss across species. Collectively, our findings showed that OCRL deficiency leads to reduced mitochondrial activity, decreased mtDNA levels, reduced mitochondrial content (TOM20), and increased oxidative stress. We further showed that OCRL- deficient neural cells exhibited an altered balance of neuronal versus astrocytic differentiation, rather than a defect in neurogenesis. Additionally, we observed impaired Sonic Hedgehog (Shh) signaling and ciliary homeostasis. Thus, we propose that mitochondrial dysfunction-induced oxidative stress acts as a central mediator linking OCRL loss to altered cell fate and disrupted Shh signaling.

