## supplement materials for "Defective Neuronal Differentiation in Lowe Syndrome is Associated with Mitochondrial Dysfunction and Impaired Cilia-related Sonic Hedgehog Signaling"

| iPSCs-QPCR primer | primer (5’ to 3’) |
| --- | --- |
| *Foxg1-F* | CCTGCCCTGTGAGTCTTTAAG |
| *Foxg1-R* | GTTCACTTACAGTCTGGTCCC |
| *Neun-F* | GTAGAGGGACGGAAAATTGAGG |
| *Neun-R* | CATAGAATTCAGGCCCGTAGAC |
| *Brn2-F* | AAAGTAACTGTCAAATGCGCG |
| *Brn2-R* | GCTGTAGTGGTTAGACGCTG |
| *Gfap-F* | CCTCCAGCGATTCAACCTTT |
| *Gfap-R* | GAAGCTCCAAGATGAAACCAAC |
| *Co2-F* | CCCCACATTAGGCTTAAAAACAGAT |
| *Co2-R* | TATACCCCCGGTCGTGTAGCGGT |
| *Dloop-F* | TATCTTTTGGCGGTATGCACTTTTAACAGT |
| *Dloop-R* | TGATGAGATTAGTAGTATGGGAGTGG |
| *Shh-F* | CGGAGCGAGGAAGGGAAAG |
| *Shh-R* | TTGGGGATAAACTGCTTGTAGGC |
| *Gli1-F* | TCTGCCCCCATTGCCCACTTG |
| *Gli1-R* | TACATAGCCCCCAGCCCATACCTC |
| *Patch1-F* | CGGCGTTCTCAATGGGCTGGTTTT |
| *Patch1-R* | GTGGGGCTGCTGTTTCGGGTTCG |
| *Actin-F* | TCACCCACACTGTGCCCATCTACGA |
| *Actin-R* | CAGCGGAACCGCTCATTGCCAATGG |

**Supplementary Figure 1. qPCR primers for iPSCs.**

| WT and IOB mouse -QPCR primer | primer (5’ to 3’) |
| --- | --- |
| *neun-F* | GTTGCCTACCGGGGTGCACAC |
| *neun-R* | TGCTCCAGTGCCGCTCCATAAG |
| *pax6-F* | GTTCCCTGTCCTGTGGACTC |
| *pax6-R* | ACCGCCCTTGGTTAAAGTCT |
| *brn2-F* | AGAGCCCAAGGCAGAAAAGT |
| *brn2-R* | GGCGCTCTGGTTAAAGGAG |
| *gfap-F* | GGACTGAACCATGTCCTTTGTC |
| *gfap-R* | AGGCTAGCTCTATCGGTATAACCTAA |
| *Mito1-F* | CTAGAAACCCCGAAACCAAA |
| *Mito1-R* | CCAGCTATCACCAAGCTCGT |
| *Cox1-F* | TGCTAGCCGCAGGCATTACT |
| *Cox1-R* | CGGGATCAAAGAAAGTTGTGT |
| *gli1-F* | TTATGGAGCAGCCAGAGAGA |
| *gli1-R* | GAGCCCGCTTCTTTGTTAAT |
| *gli2-F* | TGAAGGATTCCTGCTCGTG |
| *gli2-R* | GAAGTTTTCCAGGACAGAACCA |
| *gli3-F* | AAGCGGTCCAAGATCAAGC |
| *gli3-R* | TGTTCCTTCCGGCTGTTC |
| *patch1-F* | TGACAAAGCCGACTACATGC |
| *patch1-R* | AGCGTACTCGATGGGCTCT |
| *gapdh-F* | CATCACTGCCACCCAGAAGACTG |
| *gapdh-* *R* | ATGCCAGTGAGCTTCCCGTTCAG |

**Supplementary Figure 2. qPCR primers for wild-type and IOB mouse.**

**Supplementary Figure 3. Genotyping processing for wild-type and IOB mouse.**

(a) Genotyping primers for wild-type and IOB mouse. (b) Genotyping processing for wild-type and IOB mouse.

**(a) Genotyping primers**

| OCRL Wild-type (250 bp) | primer (5’ to 3’) |
| --- | --- |
| GT-F | **CCCTTTTCATCTGTTAGGAGAAAT** |
| GT-R | **GCATGGTTAAACGCACTATGTGG** |
| OCRL KO (480 bp) | **primer (5’ to 3’)** |
| OCRL KO –F | **GCCCTTTGATTCTAATCCCTTTTCATC** |
| OCRL KO –R | **TCTGAGCCCAGAAAGCGAAG** |

| Inpp5b (wt 150 bp/ mut 220 bp) | primer (5’ to 3’) |
| --- | --- |
| E25-F | **TAAAGTCTGAAAATCCAAGGC** |
| E34-R | **CTCATTTCTCCTTGATTCCAA** |

| INPP5B HumanBAC (240 bp) | primer (5’ to 3’) |
| --- | --- |
| in Exon 1F | **CCACCCCACGATTGACTC** |
| in Exon 1R | **GGTGTCCCAGCCCTCAG** |

**(b) Genotyping processing**

Step 1 - OCRL WT	Step 2- OCRL KO

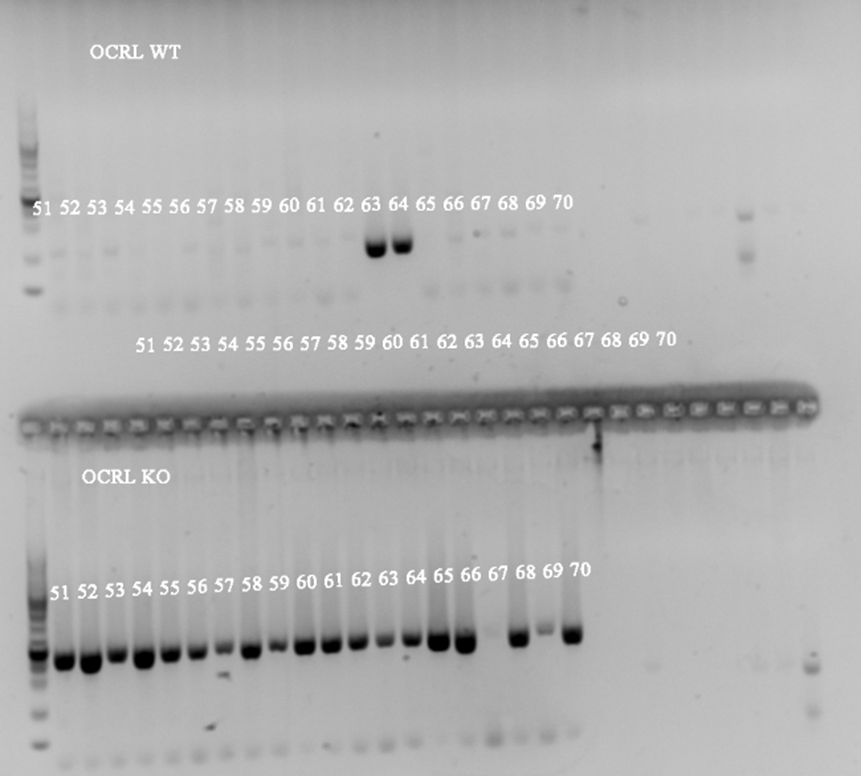

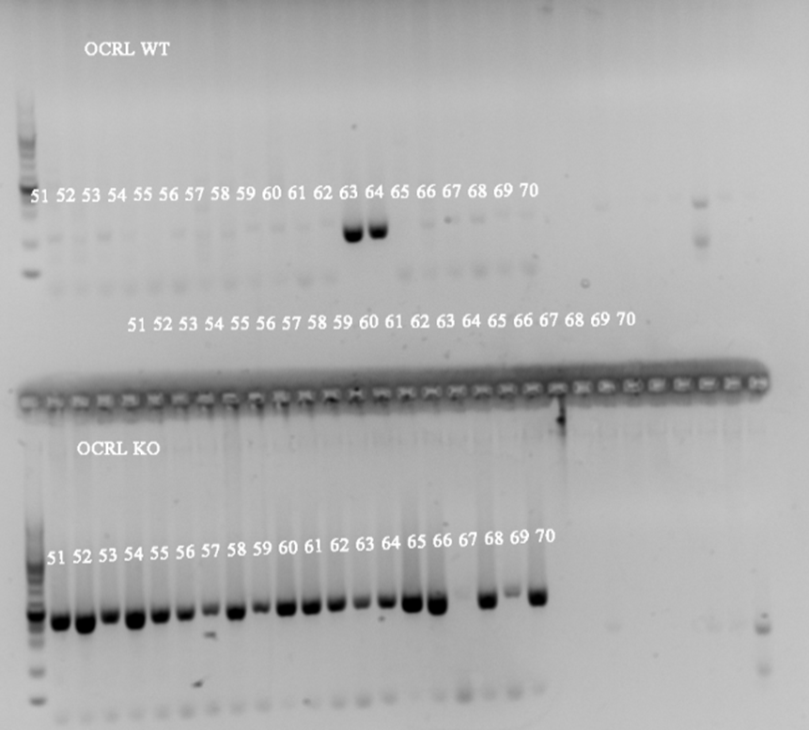

X

X

X

X

X

Step 3- Inpp5b (wt 150 bp/ mut 220 bp)	Step 4- INPP5B HumanBAC (240 bp)

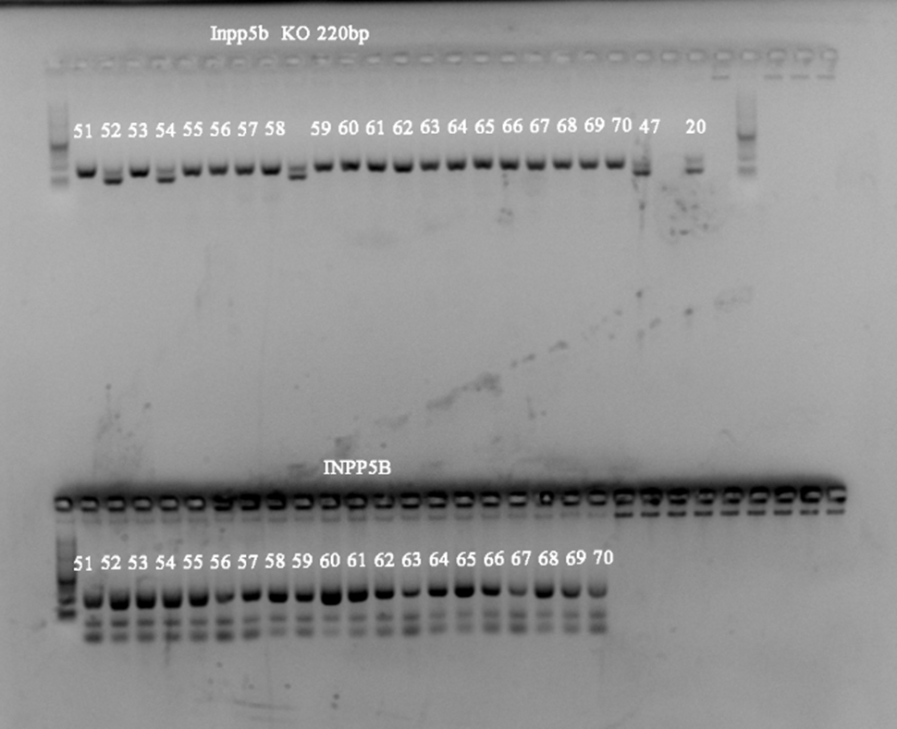

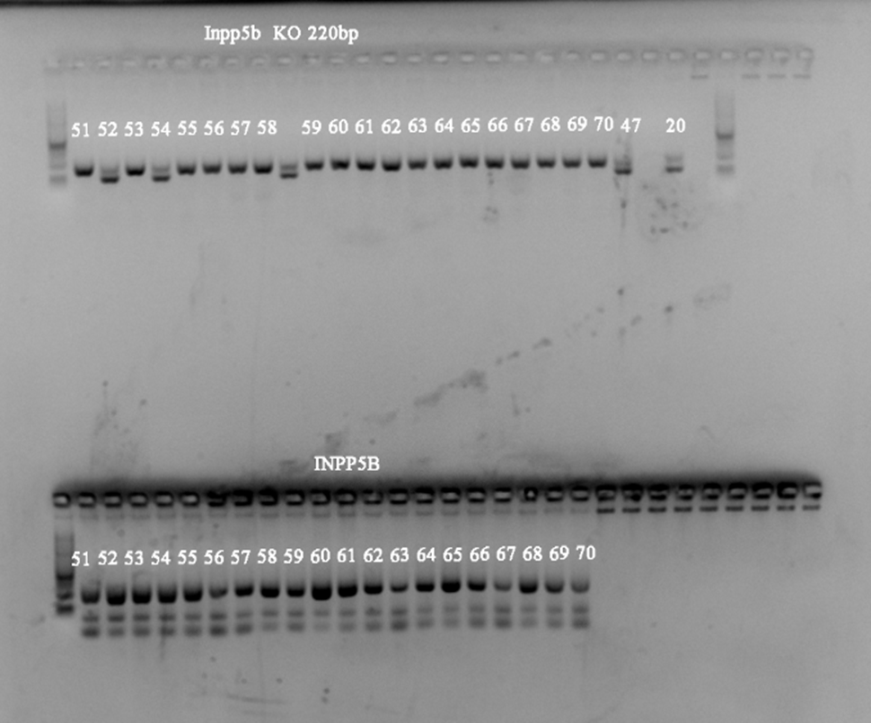

X

X

X

X

X

X

X

X

X

X

X

**Supplementary Figure 4 Validation of ocrl knockout zebrafish**

a) Two guide RNAs (gRNAs) were designed to target the ocrl locus and co-injected with Cas9 into zebrafish embryos.

**gRNA sequences:**

**gRNA1:** 5’-TCTAACAAGGACAGGAGTCTTGG-3’

**gRNA2:** 5’-TCTGCGAGGTGAACGAACACCGG-3’

**(b)** Genotyping workflow used to distinguish wild-type and ocrl-KO alleles. Genomic DNA was extracted from larvae and amplified by PCR using the following primers:

**Forward:** 5’-AGAATGTGACACTGTTTCTC-3’

**Reverse:** 5’-GTGCAGGGAAAATAAGGTGT-3’

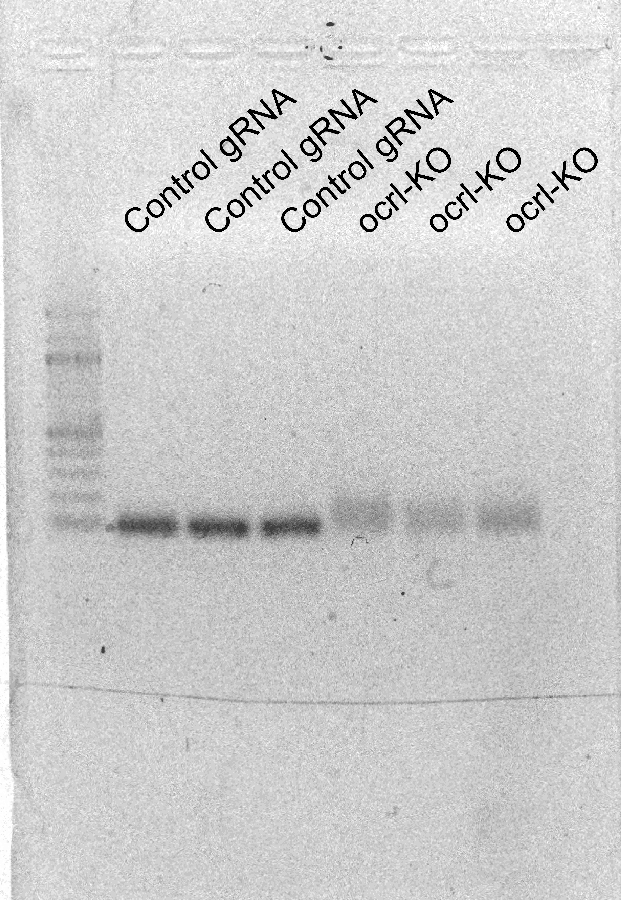
PCR products were analyzed to confirm successful genome editing at the target locus.
